# AGO104 is an RdDM effector of paramutation at the maize *b1* locus

**DOI:** 10.1101/2021.04.21.440773

**Authors:** Juliette Aubert, Fanny Bellegarde, Omar Oltehua-Lopez, Olivier Leblanc, Mario A. Arteaga-Vazquez, Robert A. Martienssen, Daniel Grimanelli

## Abstract

Paramutation is an exception among eukaryotes, in which epigenetic information is conserved through mitosis and meiosis. It has been studied for over 70 years in maize, but the mechanisms involved are largely unknown. Previously described actors of paramutation encode components of the RNA-dependent DNA-methylation (RdDM) pathway all involved in the biogenesis of 24-nt small RNAs. However, no actor of paramutation have been identified in the effector complex of RdDM. Here, through a combination of reverse genetics, immunolocalization and immunoprecipitation (siRNA-IP) we found that ARGONAUTE104 (AGO104), AGO105 and AGO119 are members of the RdDM effector complex in maize and bind siRNAs produced from the tandem repeats required for paramutation at the *b1* locus. We also showed that AGO104 is an effector of the *b1* paramutation in maize.

**Author summary:** Reprogramming of epigenetic information has been described in both plants and mammals. Here, we show that maize *ARGONAUTE (AGO) AGO104* and *AGO105/AGO119*, respectively the close homologs of *A. thaliana AGO9* and *AGO4*, are required to enable paramutation at the *b1* locus in maize. Paramutation is an epigenetic phenomenon that is stable over many generations (both mitotically and meiotically). A classic example is the *booster1* (*b1*) gene in maize, where the weakly expressed *Booster’* (*B’*) allele stably decreases the expression of the *Booster-Intense* (*B-I*) allele, and changes it into a new *B’* allele. This new *B’* allele will in turn change *B-I* into new *B’* in subsequent crosses. Previous research demonstrated that paramutation requires several proteins involved in the biosynthesis of small interfering RNAs (siRNAs) all related to the RNA-dependent DNA-methylation (RdDM) pathway. Yet, few members of the RdDM were functionally identified in maize. Here, we identify two new members of the maize RdDM pathway, and provide evidence that they are also involved in paramutation at the *b1* locus.

## Introduction

Paramutation is defined as the heritable transmission of epigenetic information that is both mitotically and meiotically stable [1–4]. There are four classical examples of paramutation in maize: the *red1 locus* (*r1*), the *plant color1* locus (*pl1*), the *pericarp color1* locus (*p1*), and the *booster1* locus (*b1*). Paramutation at the *b1* locus that encodes a transcription factor involved in the biosynthesis of anthocyanin pigments is one of the best characterized systems [5–7]. It involves the interaction between two alleles, the *BOOSTER-INTENSE* allele (*B-I*), in which *b1* is highly expressed and results in dark purple pigmentation in most mature vegetative tissues, and the *BOOSTER’* (*B’*) allele in which *b1* is weakly expressed and the plants are lightly pigmented. *B’* induces the meiotically stable *trans*-silencing of *B-I* and once *B-I* is changed into *B’* the change is permanent. Paramutation at the *b1* locus requires the presence of 7 tandem repeats (*b1TR*) located ~100 kb upstream of *b1*. The *b1TR* produce 24 nucleotide (nt) small interfering RNAs (siRNAs) through the RNA-dependent DNA Methylation (RdDM) pathway that are required for paramutation [1,5,6]. Expression of *b1TR*-siRNAs from a transgene expressing a hairpin RNA recapitulates all features of *b1* paramutation [7].

Previous studies demonstrated that paramutation has an establishment phase in developing embryos and a maintenance phase in somatic cells (Reviewed in [8]). There is evidence that the RdDM pathway is critical for both establishment and maintenance of paramutation in maize [9–12]. The RdDM pathway has been extensively described in *Arabidopsis thaliana* (Fig 1) (reviewed in [13–16]). It differs significantly in maize, with several of the catalytic subunits specific to either RNA POLYMERASE (POL) POLIV or POLV [17]. RdDM in maize is responsible for two main functions. The first one is devoted to the biogenesis of 24-nt siRNAs and the second one, called the effector complex, uses these siRNAs as guides to target chromatin and lead to DNA methylation. In the first step, POL IV transcripts are immediately converted into double-stranded RNAs (dsRNAs) by MEDIATOR OF PARAMUTATION1 (MOP1), the homolog of *A. thaliana* RNA-DEPENDENT RNA POLYMERASE (RDR) RDR2. DICER-LIKE3a (DCL3a) then slices these dsRNAs into 24-nt siRNAs [18,19]. The effector complex induces DNA methylation at either CG, CHG or CHH sites (where H=A, T, or C). In *A. thaliana*, it initiates with AGO4/6/9 [20,21], that guide the siRNAs to the transcripts generated by POL V. POL V also interacts with DEFECTIVE IN RNA DIRECTED DNA METHYLATION 1 (DRD1), DEFECTIVE IN MERISTEM SILENCING 3 (DMS3), and RNA-DIRECTED DNA METHYLATION 1 (RDM1), to produce long noncoding scaffold transcripts. This complex then recruits two histone methyltransferases of the SUVH family, SUVH2 and SUVH9, that can bind DNA but have lost catalytic activity [15]. The AGO4/6/9 complex then partners with DOMAINS REARRANGED METHYLTRANSFERASE (DRM), DRM2 and DRM1, to enable DNA methylation in all sequence contexts [13,16,17]. By sequence similarity, putative homologues of AtAGO4/6/9 have been proposed in maize. *ZmAGO104* is the closest homologue of *AtAGO9* and both *ZmAGO105* and *ZmAGO119* are the most probable homologues for *AtAGO4* [22]. To date, RdDM members found to affect paramutation in maize include MOP1 [9,23] and two REQUIRED TO MAINTAIN REPRESSION (RMR), RMR6/MOP3 that encodes the largest subunit of POL IV [9–12] and RMR7/MOP2 that encodes a subunit shared between POL IV and POLV [24–26]. These proteins are essential to maintain the paramutation states, especially MOP1 as illustrated by the dark purple phenotype in *mop1* mutant progenies [23]. Interestingly, although RMR7/MOP2 is also involved in POL V machinery, no RdDM actor from the effector complex was identified (Fig 1).

**Fig 1.**
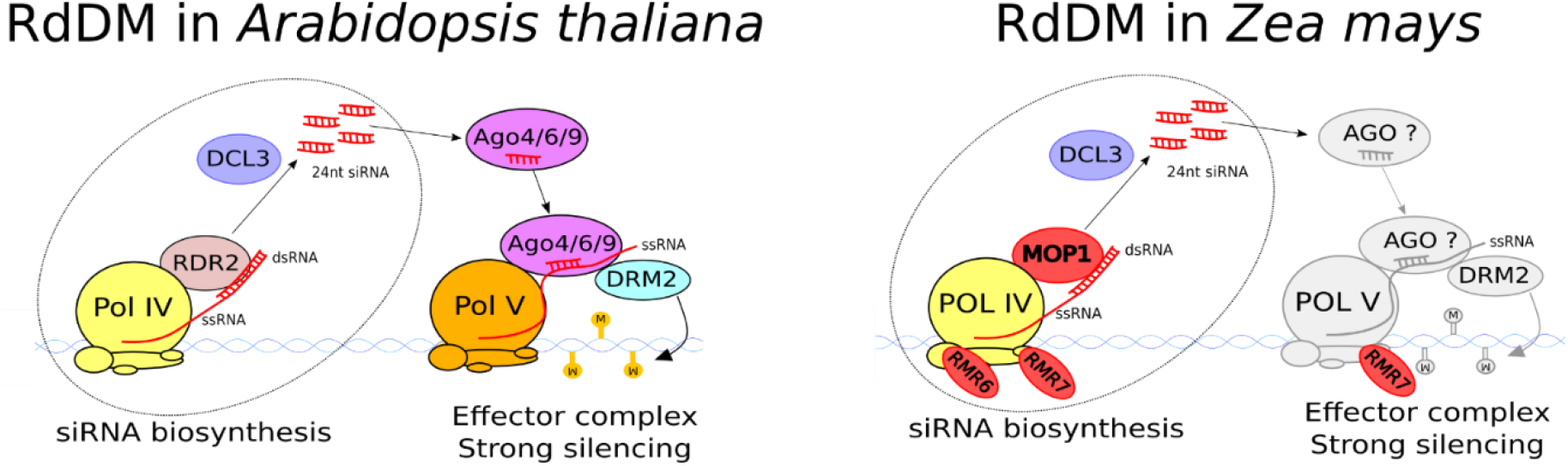
Illustration of the actors of the two steps of the RNA directed DNA Methylation (RdDM) in *Arabidopsis thaliana* and *Zea mays*. The red font color with MOP1, RMR6 and RMR7 are involved in paramutation by performing siRNA biosynthesis. RMR7 is a subunit of both POLIV and POLV. Grey font color show proteins that were not identified in maize yet but added here as hypothetic effectors by homology with *A. thaliana*.

The goal of this work was to determine whether AGO104 and/or AGO105 are involved in paramutation as part of the RdDM effector complex. We used a reverse-genetics approach to search for paramutation-associated phenotypes in *ago104* and *ago105* mutants. A novel intermediate plant pigmentation was identified in the progeny of *ago104* mutant. Using immunolocalization and immunoprecipitation, we showed that AGO104 and AGO105 have a similar function and localization to that of their homologs in *A. thaliana*. We sequenced the small RNAs bound by AGO104 and AGO105/AGO119 and showed that they target the *b1TR* repeats. Taken together, this data indicate that we identified AGO104 and AGO105/AGO119 as new members of the RdDM effector complex in maize, and we showed that AGO104 is also involved in paramutation at the *b1* locus. This research provides a deeper understanding of the establishment of paramutation as well as new insights into the role of RdDM in maize.

## Results

### Reverse genetics shows an intermediate phenotype in *ago104-5*

To study the involvement of AGO104 and AGO105 in paramutation, we selected two mutator-induced alleles, respectively *ago104-5*, previously characterized by [22] as a dominant allele creating defects during female meiosis and apomixis-like phenotypes, and the uncharacterized *ago105-1* (S1a Fig). Both mutations were backcrossed three times to the B73 inbred line, that is neutral for paramutation at the *b1* locus as it carries a *b* allele with a single tandem repeat [27].

We performed three successive crosses to generate a paramutagenic population of plants combining *mop1-1* and either *ago104-5* or *ago105-1* mutant alleles (S1b Fig). To do so, we firstly crossed recessive homozygous *mop1-1* mutant (dark purple *B’* plants) to heterozygous *mop1-1* mutant (lightly pigmented *B’* plants). The resulting progeny were only *B’* plants (either homozygous or heterozygous for *mop1-1*). After validation by genotyping, homozygous *mop1-1* mutants (dark purple *B’* plants) were crossed with either homozygous *ago104-5* or *ago105-1* mutants (i. e. green plants that harbor a *b* allele neutral to paramutation). All the resulting progeny were double heterozygous mutant for *mop1-1* and either *ago104-5* or *ago105-1* (*B’/b* plants). We evaluated plant pigmentation in these double mutants for our reverse genetic screening. All observed phenotypes have to be stable through meiosis in order to be relevant to paramutation. Consequently, our 3^rd^ crossing scheme consisted in a backcross between the double heterozygous mutants (green *B’/b* plants) and homozygous *mop1-1* plants (dark purple *B’* plants). The resulting progeny could be either single *mop1-1* mutant, or double *ago/mop1-1* mutants, with either *b* or *B’* allele. After validation by genotyping, we only conserved plants with *B’* allele that were heterozygous for *mop1-1*, or double heterozygous for *mop1-1/ago*, and evaluated plant pigmentation.

The involvement of AGO104 and AGO105 in paramutation was studied by evaluating the phenotypes at 46 days post-seeding (dps) of the previous crosses. We hypothesized that plants with disrupted paramutation would exhibit the dark purple phenotype without being homozygous for *mop1-1*. As expected for the control plants, all the homozygous *mop1-1* plants were dark purple at 46 dps, while all wild type plants were lightly pigmented. Interestingly, a new phenotype emerged in the progeny of the backcrossed double heterozygous *mop1/mop1-1;Ago104-5/ago104-5* mutants. Seven of them showed typical lightly pigmented tissues while 16 plants exhibited a previously unseen phenotype characterized by intermediate pigmentation levels, suggesting that anthocyanin production was increased compared to that of *B*’ but could not achieve the typical dark purple phenotype of *B-I* plants at 46 dps (Fig 2). We quantified the levels of pigmentation from the lightly pigmented, intermediate and dark purple phenotypes at 46 dps using picture processing on ImageJ. We measured pixel color from husk tissues of the 3 phenotypes, and both ANOVA (p-value = 0,00385) and Tukey’s ‘Honest Significant Difference’ test (p-value between 0,003 and 0,02) indicated a significant color difference between the 3 phenotypes. The pigmentation turns darker over time, and reaches levels at 56 dps similar to that observed in *mop1-1/mop1-1* (S2 Fig). Interestingly, this new phenotype happened only when the *ago104-5* allele was present in at least one parent, indicating that the parental genotype might influence the progeny’s phenotype. The partial reversion of the paramutation phenotype associated with *ago104-5* allele suggests that AGO104 is an effector of paramutation. On the other hand, the *ago105-1* mutant caused no phenotype and thus our crossing scheme did not allow evaluating its role in paramutation. Because of high sequence similarity between *ago105* and *ago119*, we crossed *ago105-1* and *ago119-1* mutants, but the presence of both mutations was systematically lethal in the progeny.

**Fig 2.**
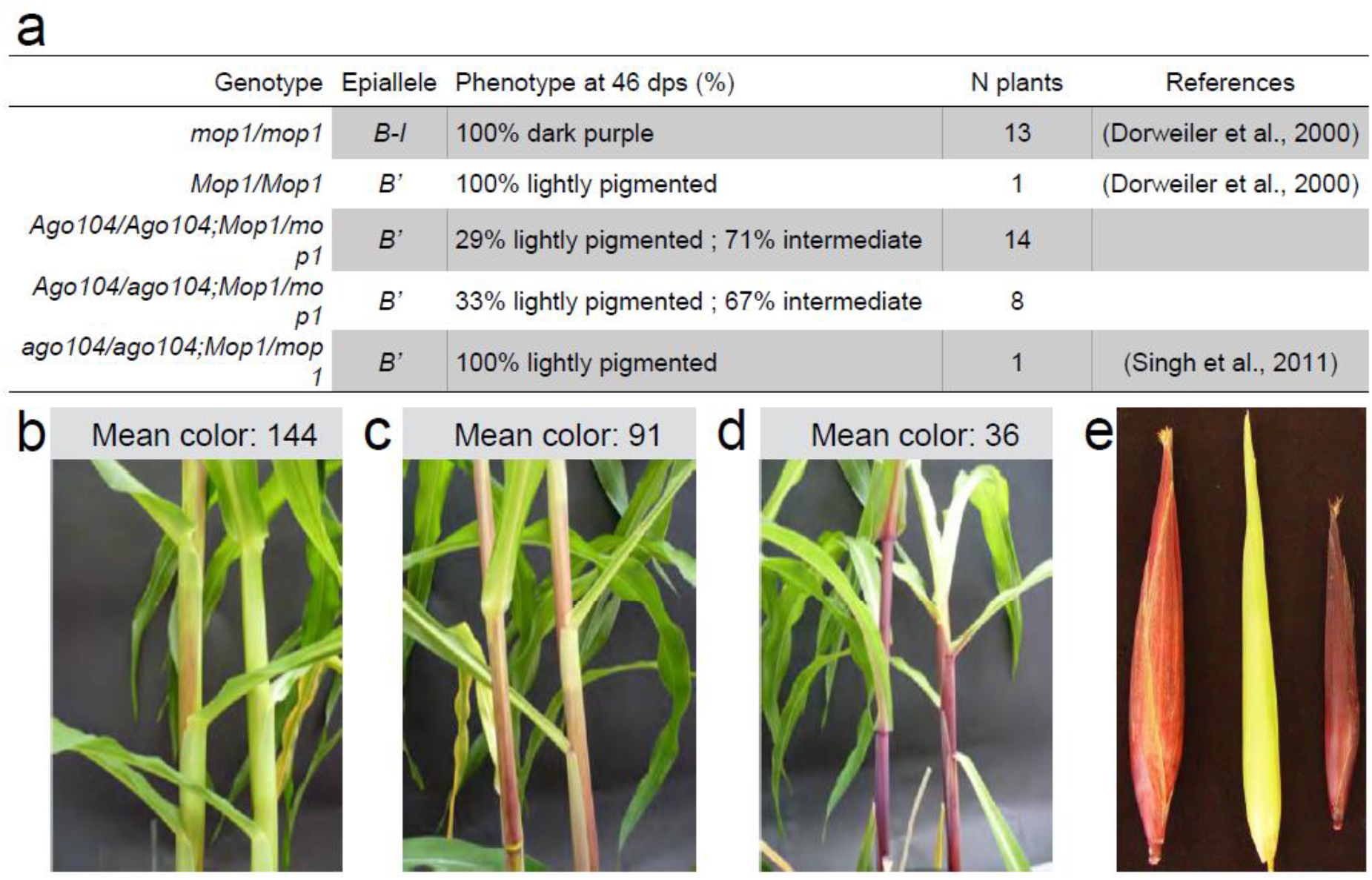
Occurrence of the intermediate pigmentation in the paramutagenic population. (a) Percentage of plants with either dark purple, light purple or intermediate phenotype for each genotype at 46 dps. The total number of plants obtained for each genotype is indicated. (b) Stem with a lightly pigmented phenotype. (c) Stem with an intermediate phenotype. (d) Stem with a fully pigmented dark purple phenotype. (e) Ears with intermediate, light purple and dark purple phenotype (from left to right). Mean color indicates the average pixel color from 2 different husk of each phenotype. It is evaluated from 0 (black pixel) to 255 (white pixel). Results from the 3 phenotypes are statistically different from each other (ANOVA p-value = 0,00385 and Tukey’s ‘Honest Significant Difference’ p-value < 0,02).

### AGO104 and AGO105/AGO119 bind siRNAs in embryonic cells

To establish the temporality of biogenesis of siRNAs in maize, we extracted small RNAs from mature and immature ears, and pollen in a heterozygous *mop1-1* (*B’* epiallele) plant and in a homozygous *mop1-1* (*B-I* epiallele) plant. We used stem-loop RT-PCR to amplify three selected siRNAs of 24-nt, namely R3, S3 and S4. R3 siRNAs are produced from transposons in a RdDM-dependent manner [28] and were used as positive control. S3 and S4 siRNAs are transcribed from the *b1TRs* and are involved in paramutation [7]. As expected, R3 was expressed in the *B’* plants, but not in the *mop1-1* RdDM mutant. In contrast, *b1TRs’* siRNAs (S3 and S4) were detected in tissues from both *B’* and *mop1-1* plants (Fig 3a), suggesting that a second pathway produces the paramutation-linked siRNAs in reproductive tissues.

**Fig 3.**
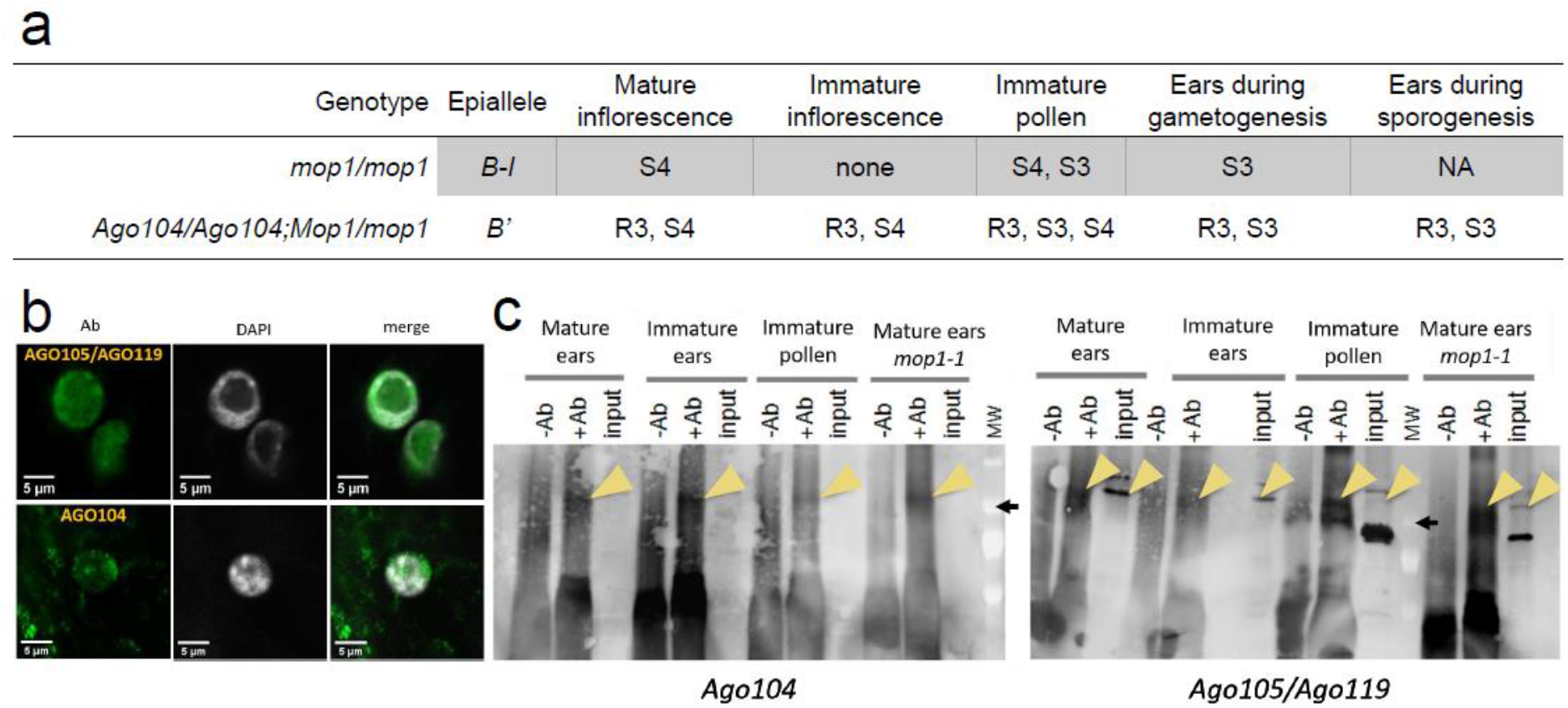
Location of *b1TR* siRNAs and AGO104/AGO105/AGO119 in reproductive tissues. (a) Results of stem-loop RT-PCR in five reproductive tissues for R3, S3 and S4 siRNAs. siRNAs class is indicated when detected. Results were identical in *B’* plants with light and intermediate pigmentation phenotypes. NA: no data available. (b) Fluorescence of AGO105/AGO119 and AGO104 in nuclei of B73 embryonic cells. (c) Immunoprecipitation of AGO104 and AGO105/AGO119 in four reproductive tissues. Yellow arrowheads indicate the location of the expected band. +Ab and -Ab are the IP samples treated with and without antibodies, respectively. Input is the sample that did not undergo IP. MW is the molecular weight, the black arrow indicates 100 kD.

To be relevant to our study, these siRNAs must be expressed in the same tissues and at the same time as AGO104 or AGO105. Therefore, we conducted an immunolocalization experiment using an anti-AGO104 antibody to determine the cellular location of AGO104 in young embryos of the B73 inbred line. We found that AGO104 was expressed in the cytoplasm of embryonic cells (Fig 3b). This profile is similar to that of AGO9 in *A. thaliana* [20]. Similarly, using an antibody against a common peptide of AGO105 and AGO119, we found that both proteins are expressed specifically in the nucleus of the embryo’s cells (Fig 3b). This data strengthens the hypothesis that *ago104* is an orthologue of *AtAGO9*, and *ago105/ago119* are orthologues of *AtAGO4*. We later performed an immunoprecipitation (IP) of AGO104 and AGO105/AGO119 in mature and immature ears and pollen. The results showed a strong expression of these proteins in immature reproductive tissues, mostly in female reproductive organs (Fig 3c). Finally, we wanted to verify whether the *mop1-1* mutation altered the AGO protein repertoire in maize and we conducted IPs using AGO104 and AGO105/AGO119 as baits in *mop1-1* mutant. The three AGOs were detected in reproductive tissues of *mop1-1* mutant suggesting that, contrary to that observed in *A. thaliana* [21], reduced levels of small RNAs in maize do not alter the integrity of ARGONAUTE proteins.

We then extracted the small RNAs from the IPs mentioned above. We hypothesized that siRNAs that are carried by AGO104, AGO105 or AGO119 should be correctly amplified and visible on a migration gel after a stem-loop RT-PCR. As expected, the control R3 siRNA (RdDM-dependent) was amplified and visible on the migration gel (S3b Fig). Therefore, our protocol allows to extract and identify the siRNAs loaded in AGO proteins. Interestingly, R3 siRNAs extracted from the IPs of mature ears were less abundant than those detected in immature ears. In contrast, we were not able to visualize S3 siRNAs involved in paramutation in both mature or immature ears (S3b Fig). However, when extracted directly from mature and immature ears, S3 siRNAs could be detected. We can draw two hypotheses from this result: either AGOs do not bind S3 siRNAs involved in paramutation, or our experiment using stem-loop RT-PCR is not sensitive enough to visualize it.

### AGO104 and AGO105/AGO119 bind 24-nt siRNAs involved in paramutation

We then sequenced the small RNAs recovered from immature ears of AGO104 IPs in plants producing normal and reduced amounts of 24-nt siRNAs. The normal 24-nt siRNA production is represented by B73 plants (*b* allele) and heterozygous *mop1-1* mutant (*B’* epiallele), and the reduced 24-nt siRNA production is represented by homozygous *mop1-1* mutant (*B-I* epiallele) and homozygous *ago104-5* mutant (*b* allele). We first evaluated the expression level for the three studied siRNAs in each genotype. Interestingly, we identified R3 siRNAs in all four genetic backgrounds, which can be explained by a weak sensitivity of the previous stem-loop RT-PCR experiments. Moreover, we identified S3 and S4 siRNAs in none of the genetic backgrounds (S3c Fig). This means that AGO104 does not bind the S3 and S4 sRNA in immature ears. Next, we aligned AGO104-associated small RNAs onto the B73 reference genome, and all genotypes displayed a very similar chromosome-scale coverage (S4 Fig). We evaluated the size of the reads and mapped them to a 100kb where we replaced the *b1* promoting sequences of B73 genome by the *b1TR’s* repeats [5] (accession AF483657) (Fig 4). AGO104 from plants producing normal amounts of 24-nt siRNA (heterozygous *mop1-1* mutant, B’ epiallele) carried mostly 24-nt small RNAs, which mapped to the *b1TR’s* region. This result indicates that AGO104 from heterozygous *mop1-1* (*B’* epiallele) binds the 24-nt siRNAs associated with paramutation. On the other hand, AGO104 from homozygous *mop1-1* mutant (*B-I* epiallele) carried mostly 22-nt small RNAs, which did not map to the *b1TR’s*. This indicates that in homozygous *mop1-1* plants (*B-I* epiallele), AGO104 does not bind *b1TR* associated 24-nt siRNAs. It has been shown that the *b* allele produces the same *b1TR* siRNAs as *B-I* allele [7]. Therefore, small RNAs from AGO104 of the *ago104-5* mutant (*b* allele) were also aligned on the B73 sequence with the 7 tandem repeats. Interestingly, AGO104’s small RNAs from *ago104-5* mutant display a profile similar to that of homozygous *mop1-1* plants (*B-I* epiallele). Taken together, these results indicate that AGO104 binds *b1TR* associated 24-nt siRNAs, and that its involvement is crucial to enable paramutation.

**Fig 4.**
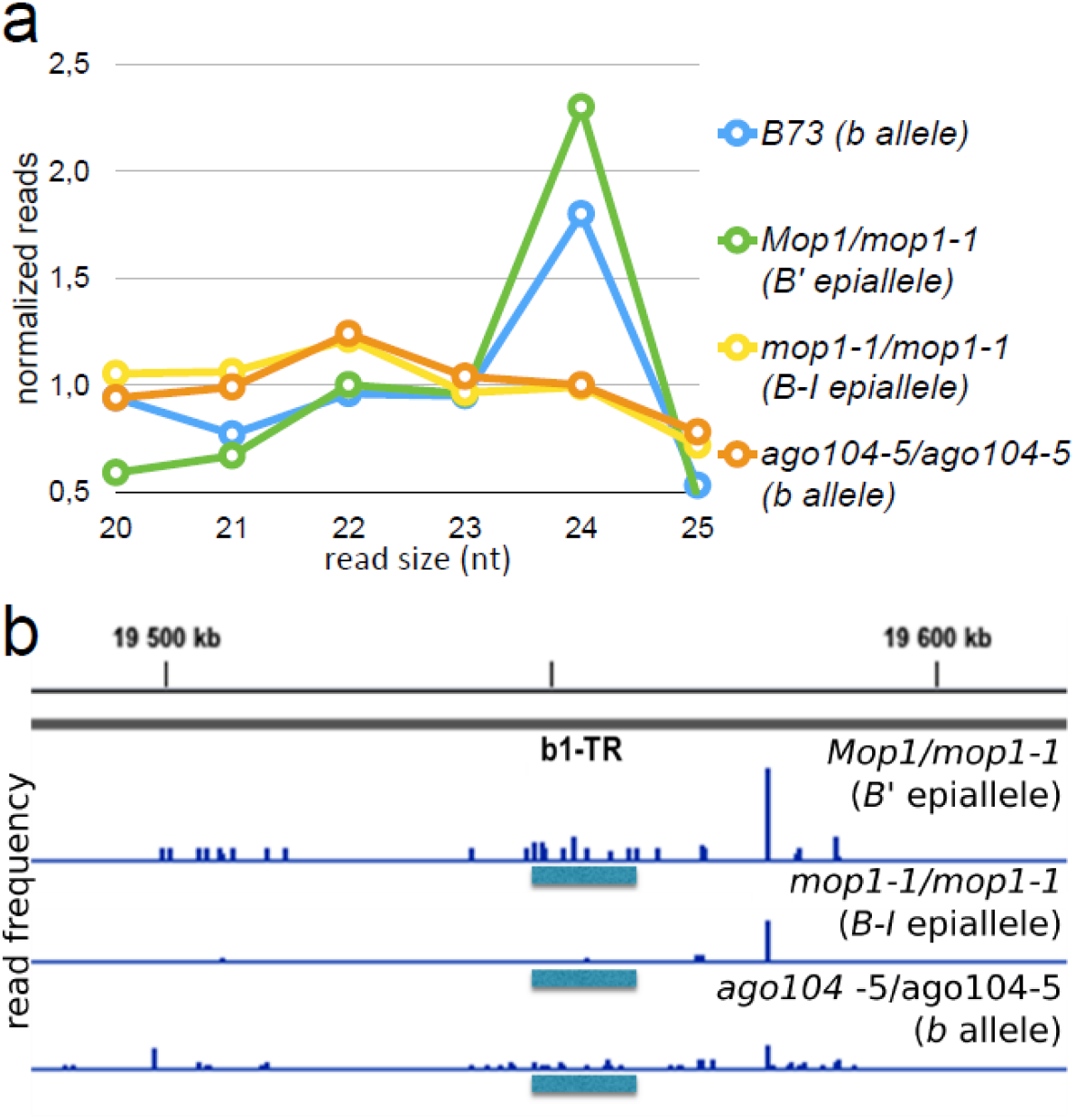
Distribution of AGO104’s small RNAs in *b* allele (B73 inbred and *ago104-5* mutant), and in *B’* (*Mop1-1* heterozygous) and *B-I* epialleles (*mop1-1* homozygous). (a) Size distribution of reads normalized to 1. (b) Reads distribution within the 100 kb region centered on *b1TR*. Blue horizontal rectangles indicate the *b1TR* location. Vertical blue bars indicate read frequency.

Finally, we verified whether AGO105/AGO119, like AGO104, binds paramutation-associated small RNAs. We sequenced the small RNA-IPs of AGO105/AGO119 and AGO104 that we generated from immature ears of wild type B73 plants (*b* allele). We also downloaded small RNAs extracted directly from B73 young ears [29] (accession GSM918110). We identified a 8-kb sequence that span the *b1*-single repeat of B73 and its upstream sequence in the B73 genome. The three small RNA datasets were mapped to the 8-kb sequence, and they covered most of the *b1*-single repeat (Fig 5). Small RNAs produced by B73 (*b* allele) at the *b1*-single repeat are identical to those produced by *B-I* at the *b1TR* locus [7]. Therefore, we can conclude that both AGO104 and AGO105/AGO119 bind *b1TR* small RNAs, and are involved in paramutation. Interestingly, the small RNAs extracted from AGO104 and those extracted from AGO105/AGO119 are quite similar: they map to the same locations on the *b1*-single repeat, and the first base preference tends to favor adenine and disadvantage thymine in both AGO104 and AGO105/AGO119 (S5 Fig). This might indicate a similar involvement in paramutation. The evidence from this study suggests that AGO105 and/or AGO119 are involved in paramutation.

**Fig 5.**
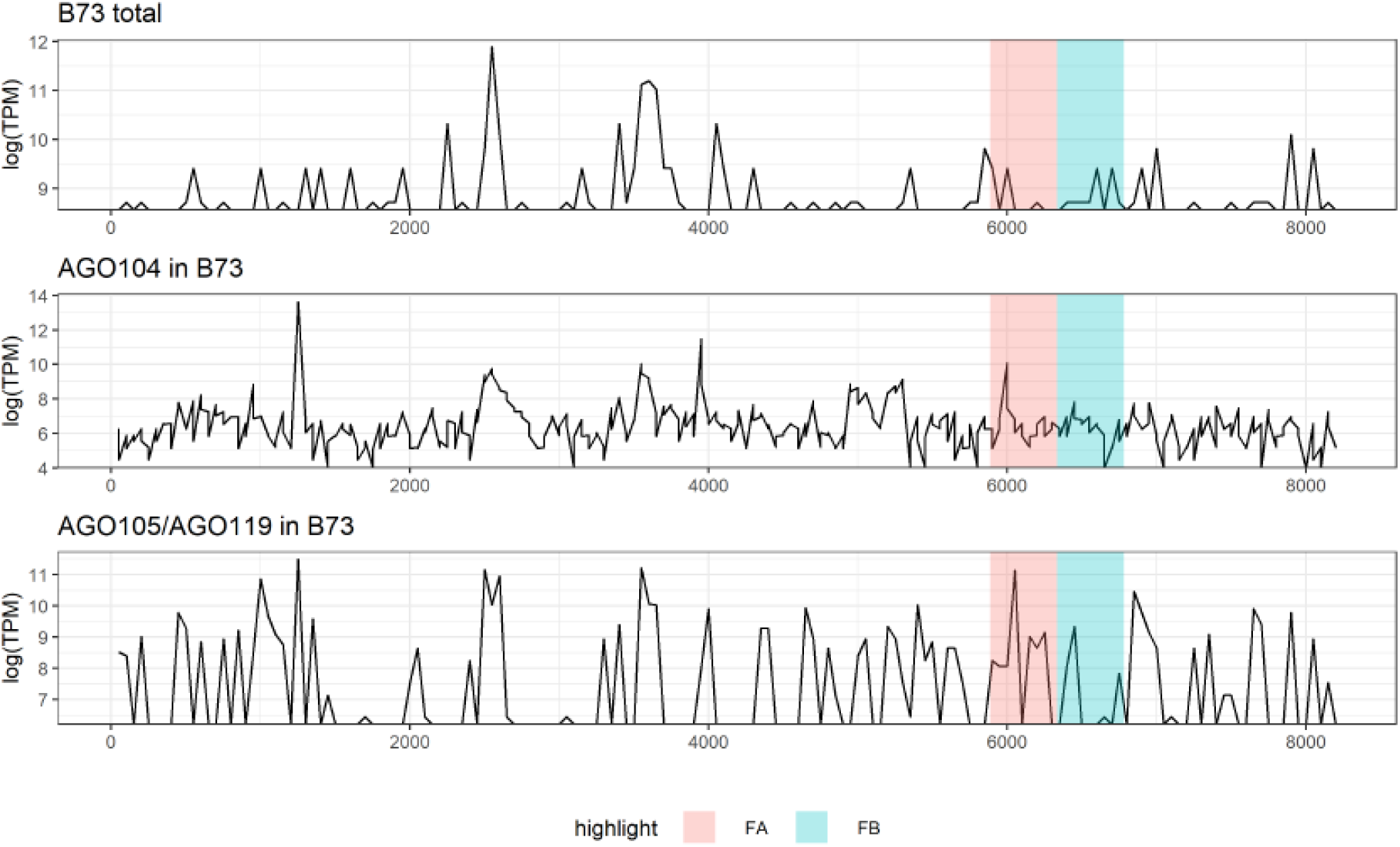
Reads distribution of small RNAs at B73 single repeat and its promotor sequence. Reads marked as “B73 total” are small RNAs extracted from B73 young ears ([29] accession GSM918110). Reads marked as “AGO104 in B73” and “AGO105/AGO119 in B73” are small RNAs respectively extracted from AGO104 and AGO105/AGO119 of B73 young ears. FA (pink box) and FB (blue box) are the two halves of the B73 single-repeat, as proposed by [32]. x-axis is in base pairs.

## Discussion

The reverse genetic screening performed on *ago104-5* and *ago105-1* mutants helps understanding their involvement in paramutation. Paramutation at the *b1* locus involves the *B-I* and *B’* epialleles, respectively associated with intense and weak plant pigmentations [30]. Here, we unveiled an intermediate plant pigmentation phenotype appearing in the progeny of *ago104-5* mutants with the *B’* allele, which turns darker over time. Previous description of *mop2* mutant also reported an evolution of pigmentation over time, but never rising up to the levels of homozygous *mop1-1* mutant [25]. This suggests that the *ago104-5* mutation disrupted paramutation in its progeny at the *b1* locus. Two conclusions can be drawn from these results. First, AGO104 is an effector of paramutation at the *b1* locus. Second, the parental genotype of *ago104-5* mutants influenced their progeny’s phenotype, with the interaction taking place in reproductive tissues. In accordance with our results of IP and immunolocalization, previous studies have demonstrated that AGO104 is located exclusively in reproductive tissues, where paramutation is established. These tissues include female and male meiocytes, egg cells, but not the gametic precursors [22]. Interestingly, *b1* is expressed in somatic tissues only [7], where maintenance of paramutation takes place, and where AGO104 is never expressed. Hence, AGO104 is probably involved in the establishment rather than the maintenance of paramutation. On the other hand, the *ago105-1* mutation did not display any phenotype that was differing from the control groups. This result may be explained by the fact that previous studies identified AGO119 as closely related to AGO105 [22]. Therefore, AGO119 might complement mutations in *Ago105* and prevent the establishment of new phenotypes. The double *ago105/ago119* mutants were lethal, which prevented further analysis on that hypothesis. Our reverse genetic screenings enabled us to identify AGO104 as an effector of paramutation at the *b1* locus.

Understanding the similarities between maize and *A. thaliana* AGO proteins is a first step to understand their function in maize. In *A. thaliana*, AGO4/6/9 have closely related sequences, closely related functions, and interact with small RNAs of the same size [21]. Within the AGO family of Arabidopsis, they belong to the same clade [31]. Maize *ago104, ago105* and *ago119* are close homologs of *AtAGO4/6/9*. Our results show that *Zm*AGO104 is present in the cytoplasm of embryos, just like *At*AGO9 [20]. Similarly, *Zm*AGO105 and *Zm*AGO119 are located in the nucleus of embryos, like *At*AGO4. This different localization in *A. thaliana’s* embryos does not prevent the involvement of both *At*AGO4 and *At*AGO9 in gamete formation [20,21]. Therefore, an involvement of AGO104, AGO105 and AGO119 in maize RdDM can be expected despite their different localization in embryonic cells. Based on our results of immunolocalization and on sequence similarities previously reported [22], we argue that maize AGO104 and AGO105/AGO119 are orthologs of *At*AGO9 and *At*AGO4, respectively. In support of this claim, as their orthologs *At*AGO9 and *At*AGO4 [21], maize AGO104, AGO105 and AGO119 bind preferentially 24-nt small RNAs. Some of these 24-nt small RNAs extracted from AGO104 and AGO105/AGO119 mapped to the *b1TRs* involved in paramutation. Prior studies have emphasized the importance of the first half of the *b1TR* sequence for the paramutagenicity of the *b1* locus [32]. Interestingly, AGO104 and AGO105/AGO119 load *b1TR*’s siRNAs that map to this first half of the repeats, even though *ago105-1* mutants did not show any phenotype. This might indicate a hierarchy in the requirement of these AGOs in paramutation, and a complementation of *ago105-1* by other AGOs, like AGO119.

To expand our comparison between the AGOs of maize and *A. thaliana*, we considered the degradation process of the *At*AGOs. In the *rdr2* mutant in *A. thaliana (mop1* in maize), the levels of 24-nt siRNAs are low, and the AGO4s, AGO6s and AGO9s are degraded [21]. Our quantification of AGO104 and AGO105/AGO119 in *mop1-1* mutant of maize showed that the decrease in 24-nt siRNAs production does not influence the stability of the AGO104 and AGO105/AGO119 proteins. This suggests that either the degradation mechanisms are different in maize or there is a pathway capable of rescuing AGO104 and AGO105/AGO119 independently of MOP1-dependent siRNAs. As previously shown in *A. thaliana*, there is more than one pathway that produces 24-nt siRNAs. Within the highly conserved RdDM mechanism, there are “canonical” and “alternative” pathways that enable the synthesis of 24-nt siRNAs without the involvement of *RDR2* [33]. The same happens in maize, which can create some 24-nt siRNAs without the involvement of MOP1 [34,35]. It is therefore logical that we could amplify paramutation-linked 24-nt siRNAs in homozygous *mop1-1* mutant (Fig 3a). However, the production of 24-nt small RNAs in the *mop1-1* mutant is partially replaced by 22-nt small RNAs [34]. This supports our results in which AGO104 proteins in homozygous *mop1-1* mutant did not carry S3 and S4 siRNAs (S3c Fig), and they carried more 22-nt small RNAs than 24-nt small RNAs (Fig 4a). A possible explanation for this might be that the 22-nt small RNAs in *mop1-1* mutant contribute to rescue AGO proteins, but they do not mediate paramutation at the *b1* locus.

This study has identified one new actor of paramutation in maize, and has shown that more ARGONAUTE proteins are involved, through a reverse genetic approach, and by sequencing small RNAs loaded onto AGO proteins. These experiments confirmed that AGO104 is involved in paramutation in maize and binds paramutation-associated siRNAs. It is involved in the establishment of paramutation in the reproductive tissues of maize through the effector complex of RdDM. Although no phenotype was associated to the *ago105-1* mutation, AGO105 and/or AGO119 were found to be involved in RdDM and to bind paramutation-associated siRNAs as well. While more actors need to be identified to complete our knowledge of the pathways involved in paramutation, our findings shed new light on the mechanisms mediating both the establishment and the transmission of paramutation in maize.

## Materials and methods

### Plant material

The *ago105-1* mutant is available at the Maize Genetics Cooperation Stock Center under reference UFMu-05281. B73 inbred line was provided by the Maize Genetics Cooperation Stock Center. The Trait Utility System for Corn (TUSC) at Pioneer Hi-Breed provided *ago104-5* stocks and V.L. Chandler (University of Arizona, Tucson, AZ, USA) provided the *mop1-1* mutants. Plants were grown in a greenhouse at the French National Research Institute for Sustainable Development in Montpellier, France, with 14 hours day light (26°C during the day, 20°C at night). For all these plants, pollen, immature and mature ears were collected and immediately snap frozen in liquid nitrogen and stored at −80°C before use.

### Immunolocalization

Fertilized ovaries from B73 plants were collected 3 days after pollination (DAP) and sliced using a Vibratom (Leica VT1000E) to create 200 to 225 μm sections. They were left 2 hours in fixating solution (4% paraformaldehyde, PBS 1X, 1% Tween 20, 0.1 mM PMSF) and washed 3 times in PBS (Phosphate Buffered Saline). Samples were then digested for 15min at room temperature using an enzymatic solution (1% driselase, 0.5% cellulase, 1% pectolyase, 1% BSA, all from Sigma-Aldrich), and washed 3 times in PBS. Samples were left 1 hour in permeabilizing solution (PBS 1X, 2% Tween 20, 1% BSA) in ice, and were then washed 3 times in PBS and incubated overnight at 4°C with primary antibodies (listed in S1 Table) concentrated at 1:50 for AGO104 and 1:200 for AGO105/AGO119. Samples were left 8 hours in washing solution (PBS 1X, 0,2% Tween 20) with solution renewal every 2 hours. They were incubated overnight in secondary antibody (1:200) labeled with Alexa Fluor 488, and left 6 hours in washing solution. They were then incubated 1 hour in DAPI, rinsed with PBS 1X, and mounted in ProLong Antifade Reagent (Invitrogen). Slides were sealed with nail polish and stored at −20°C. Observations were made using LEICA SPE with 405 nm (DAPI) and 488 nm (Alexa fluor 488) excitation.

### Small-RNA Immunoprecipitation

Protocols were adapted from [21] using two biological replicates per genotype. Tissues were grinded with liquid nitrogen and a Dounce homogenizer. Resulting powder was placed in a Falcon tube with 3 volumes of extraction buffer (20 mM Tris HCL pH 7.5, 5 mM MgCl_2_, 300 mM NaCl, 0.1% NP-40, 5 mM DTT, 1% protease inhibitor (Roche Tablet), 100 units/mL RNase OUT (invitrogen)). Samples were vortexed, kept on ice 30 minutes with recurrent shaking, and centrifuged 20 minutes at 4°C, 4000 rpm. Supernatants were filtered through a 0.45μm filter into a new Falcon tube, and 1 mL and aliquoted and stored at −20°C as a pre-experiment input sample. In the remaining samples, 2 mL aliquots were generated, and we added 5 μg of antibodies per gram of tissue. They were incubated 1h at 4°C on a rotation wheel. Magnetic beads (Dynabeads, Life technologies) were washed 3 times in wash buffer (20 mM Tris HCL pH 7.5, 5 mM MgCl_2_, 300 mM NaCl, 0.1% NP-40, 1% protease inhibitor (Roche Tablet), 100 units/mL RNase OUT (invitrogen)). 20 μL of washed beads were added to each sample and incubated on rotation wheel for 2 hours at 4°C. Beads from the samples were washed 3 times in wash buffer and resuspended in 500 μL. 100 μL was aliquoted and stored at −20°C for the Western blot control. Wash buffer was discarded and replaced by 250 μL of elution buffer (100 mM NaHCO3, 1% SDS, 100 units/mL RNase OUT (Invitrogen) in 0,1% DEPC water according to [36], and tubes incubated 15 minutes at 65°C with agitation. Supernatant was transferred to fresh tubes and elution was repeated once. The two eluates were finally combined. Samples were treated with 0.08 μg/μL proteinase K for 15 minutes at 50°C. RNAs were extracted following the recommendations from Applied Biosystems for TRI Reagent^®^ Solution, starting by adding 1.2 mL of TRI Reagent to the samples.

### Stem loop RT PCR

Small RNAs extracted from the RNA-IP were treated with DNase to remove a potential contamination with DNA, using the TURBO DNA-free kit (AM1907, Ambion Life technologies). In the DNA-free samples, 50 μM of stem-loop primer (listed in S2 Table), 10 mM of dNTP and nuclease-free water were added to reach 13 μL. The stem-loop reverse transcription was done following the recommendations from [37] resulting double stranded cDNA was used for PCR. 1 μL of cDNA was mixed with Red Taq 2x (Promega), and 0,25 μM of universal reverse primer (complementary to the stem loop one) and a specific forward primer, designed to match the *b1TR* siRNAs. The tubes filled with 20 μL of reaction were denatured for 2 minutes at 94°C, and went through 40 cycles of 15 seconds at 94°C and 1 minute at 60°C. Migration was done on 2% agarose gels (Lonza) with TBE 0.5X and 0.5 μg/mL BET. 100 bp Promega DNA Ladder was added, and migration was done 40 minutes at 100 volts. To make sure that the resulting bands are indeed the cDNAs from *b1TR* siRNA, the content of the gel bands were recovered using the QIAquick gel extraction kit (QIAGEN). Resulting DNA was cloned in DH5α competent cells (Invitrogen) using the pGEM-T Easy Vector Systems protocol (Promega) and an LB-ampicillin selective medium. Colonies were genotyped using the T7/SP6 primers (Promega). Plasmids from the validated colonies were isolated using the QIAprep Spin Miniprep Kit (QIAGEN) and sent for sequencing at Beckman Coulter genomics.

### Western blot

The protocols were adapted from [38]. Various types of tissues were collected: pollen, mature and immature ears from B73, and mature ears from *mop1-1* mutants. Tissues were grinded with liquid nitrogen and mixed with extraction buffer (125 mM Tris pH 8.8, 1% SDS, 10% glycerol, 10 mM EDTA, 1 mM PMSF, 1% protease inhibitor (Roche Tablet)). Samples were centrifuged 20 minutes at 4000 rpm at 4°C. Pellet were added 0.1 volume of Z buffer (125 mM Tris pH 6.8, 12% SDS, 10% glycerol, 5 mM DTT, Bromophenol Blue). Proteins in the supernatant were quantified using the Bio-Rad protein assay. 15 μg of proteins were aliquoted and 0.1 volume of Z buffer was added. The control samples from the RNA-IP were added Laemmli 4X (250 mM Tris pH 6.8, 4% SDS, 20% glycerol, 10 mM DTT, Bromophenol Blue), and incubated 15 minutes at 90°C. Migration was done at 180 volts for 45 minutes in migration buffer 1X (25 mM Tris base, 190 mM glycine, 0.1% SDS) with PageRuller plus ladder (ThermoFisher Scientific). Transfer was done on a nitrocellulose membrane (Amersham) in transfer buffer (migration buffer 1X, 20% ethanol), at 125 volts for 2 hours and a half. Membrane was then rinsed in PBST and left 1 hour in 5% non-fat milk (mixed with PBST). Milk was renewed and added 1/200 antibody (Eurogentec) against AGO104 or AGO105/AGO119 and left overnight with agitation. Membrane was washed 4 times in milk, with 5 minutes agitation every time. Milk was then added with 1/2500 HRP antibody (Invitrogen) and left 2 hours with agitation. Membrane was washed 4 times with PBST and treated as recommended by ECL plus western blotting detection system (Amersham) with a Typhoon 9400. Membrane was then washed again in PBST and left 30 minutes in Ponceau S Solution (Sigma-Aldrich) with agitation before a last water washing.

### Small RNA sequencing

Small RNAs extracted from the RNA IP were migrated on a 1.5% agarose gel. Bands corresponding to small RNAs were collected and their content was recovered using the Monarch DNA Gel Extraction kit (NEB #T1020 New England Biolab). The sRNA collected were turned into libraries using the NEBNext Multiplex Small RNA Library Prep Set (NEB #E7300S New England Biolab). The final enrichment PCR was made with 15 cycles. Samples were quantified with Qubit and Agilent Bioanalyzer using the DNA high sensitivity assays and were sequenced on a NextSeq550 machine at the CSHL Genome Center.

### Small RNA seq analysis

Sequenced datasets were cleaned using Trimmomatic (Version 0.38) with parameters 2:30:5 LEADING:3 TRAILING:3 SLIDINGWINDOW:4:15 MINLEN:15 MAXLEN:35. Reads were first mapped onto B73 RefGen_V5 of the maize genome using Bowtie 1 (Version 1.2.2) with parameters --best -k 2 −5 4 - p 10. Reads were then intersected into 0.5 Mb genome windows using bedtools coverage. For a better resolution, reads were also aligned to the *b1TRs* and their 100 kb flanks via Bowtie 1 (Version 1.2.2) with parameters -m 7 - q --strata --best -v 2. They were intersected into 50 bp genome windows using bedtools coverage.

Small RNA sequencing data were deposited in the Gene Expression Omnibus (GEO) database (https://www.ncbi.nlm.nih.gov/geo/) under the accession number GSE172479.

### Quantification of plant pigmentation

For pixel color measurement, 2 husks of plants with lightly pigmented, intermediate and dark purple phenotype were scanned. Identical squares were numerically designed in each husk to ensure an equal number of pixels, and their color was measured using ImageJ, with a scale from 0 (black pixel) to 255 (white pixel). The resulting pixel color values were averaged for each phenotype and added to Fig 2.

## Acknowledgments

D.G. received support for the H2020 MCA (REP-658900-2), and *Agence Nationale de la Recherche* grants REMETH (ANR-15-CE12-0012-03) and CHROMOBREED (ANR-18-CE92-0041). M.A.A.V. received support from the Jeunes équipes associées à l’IRD (JEAI) program (EPIMAIZE), Agropolis Fondation, CONACYT (158550 & A1-S-38383) and the Royal Society Newton Advanced Fellowship (NA150181). O.O.L. was the recipient of a graduate scholarship from CONACYT and from the Bourses d’échanges scientifiques et technologiques IRD (BEST) program. R.A.M. is supported by the Howard Hughes Medical Institute, and grants from the National Science Foundation. The authors acknowledge assistance from the Cold Spring Harbor Laboratory Shared Resources, which are funded in part by the Cancer Center Support Grant (5PP30CA045508).

## Author contributions

**Conceptualization:** Juliette Aubert, Fanny Bellegarde, Olivier Leblanc, Daniel Grimanelli

**Data curation:** Juliette Aubert, Olivier Leblanc, Daniel Grimanelli

**Formal analysis:** Juliette Aubert, Fanny Bellegarde, Olivier Leblanc

**Funding acquisition:** Robert A. Martienssen, Daniel Grimanelli

**Investigation:** Juliette Aubert, Fanny Bellegarde, Omar Oltehua-Lopez, Daniel Grimanelli

**Methodology:** Juliette Aubert, Fanny Bellegarde, Mario A. Arteaga-Vazquez, Olivier Leblanc, Daniel Grimanelli

**Visualization:** Juliette Aubert, Olivier Leblanc, Daniel Grimanelli

**Writing – original draft:** Juliette Aubert, Fanny Bellegarde, Mario A. Arteaga-Vazquez, Omar Oltehua-Lopez, Olivier Leblanc, Daniel Grimanelli

**Writing – review & editing:** Juliette Aubert, Fanny Bellegarde, Mario A. Arteaga-Vazquez, Omar Oltehua-Lopez, Olivier Leblanc, Robert A. Martienssen, Daniel Grimanelli

## Supporting information

**S1 Fig. Use of *ago104-5* and *ago105-1* mutants to create a paramutagenic population for a reverse genetic screening.**

**S2 Fig. Evolution of mutant phenotypes at 35, 46 and 56 days post-seeding (dps).**

**S3 Fig. Presence of R3, S3 and S4 siRNAs loaded into AGO104 and AGO105/AGO119 in various genetic backgrounds.**

**S4 Fig. siRNA coverage on the 10 maize chromosomes of B73 reference genome (version 5).**

**S5 Fig. First base nucleotide bias in siRNAs bound by AGO104 and AGO105/AGO119.**

**S1 Table. Antibodies characteristics**.

**S2 Table. Primer sequences used for siRNA stem loop RT-PCR.**

## Notes

### Competing Interest Statement

The authors have declared no competing interest.

