## Supplemental Figures for "AGO104 is an RdDM effector of paramutation at the maize *b1* locus"

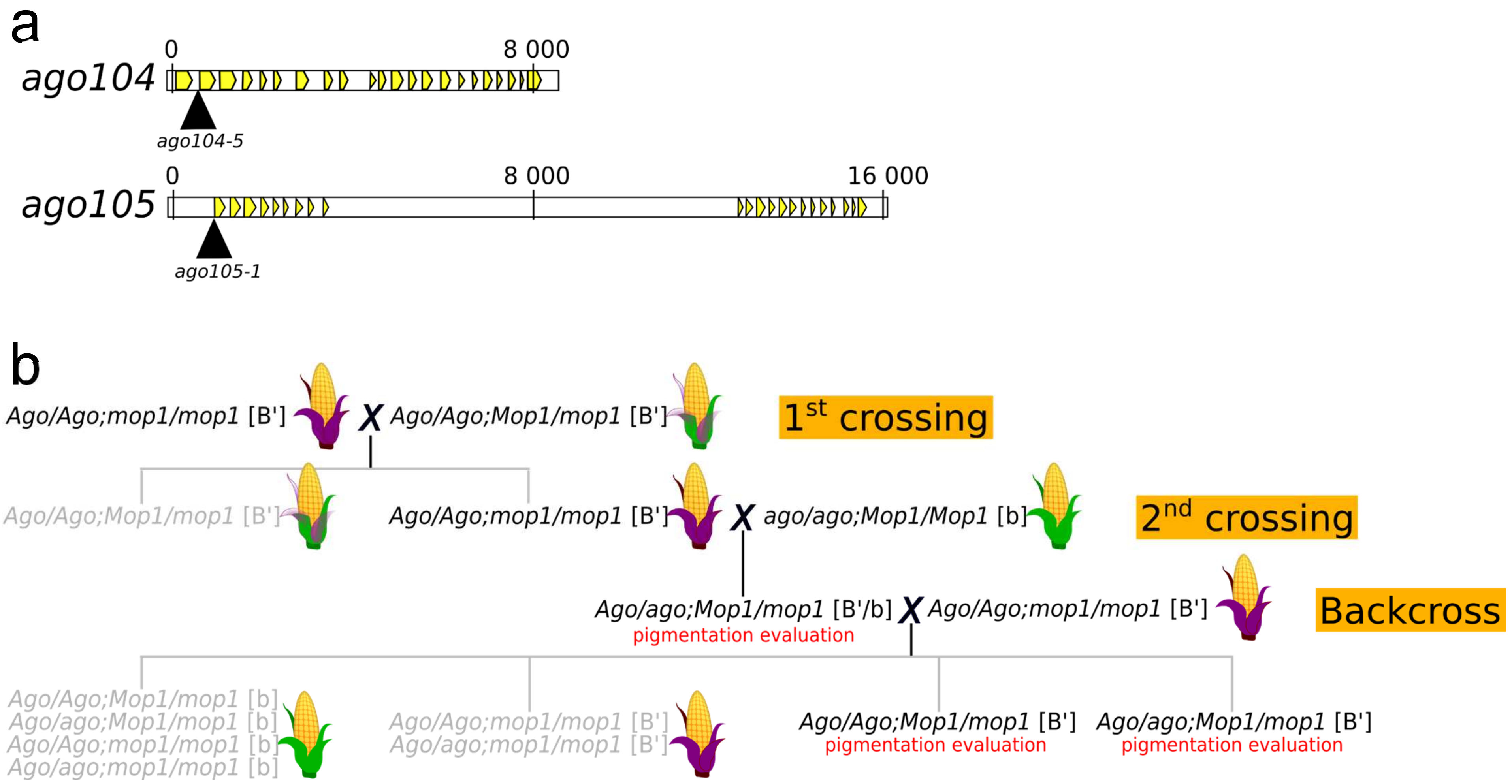


**S1 Fig. Use of *ago104-5* and *ago105-1* mutants to create a paramutagenic population for a reverse genetic screening.** (a) Schematic representation of the *ago104-5* and *ago105-1* mutants. Arrowheads indicate Mutator insertion sites. Yellow boxes indicate exons. Numbers are in base pairs, with 0 the transcription start site. (b) Scheme for the development of the paramutagenic population. This population was created twice using *ago104-5* and *ago105-1*. The *mop1* mutant is a *mop1-1* allele. Purple, lightly pigmented and green ears represent respectively dark purple, lightly pigmented and green stem phenotypes. Genotypes in grey were excluded from subsequent analyses. Alleles are indicated in square brackets.

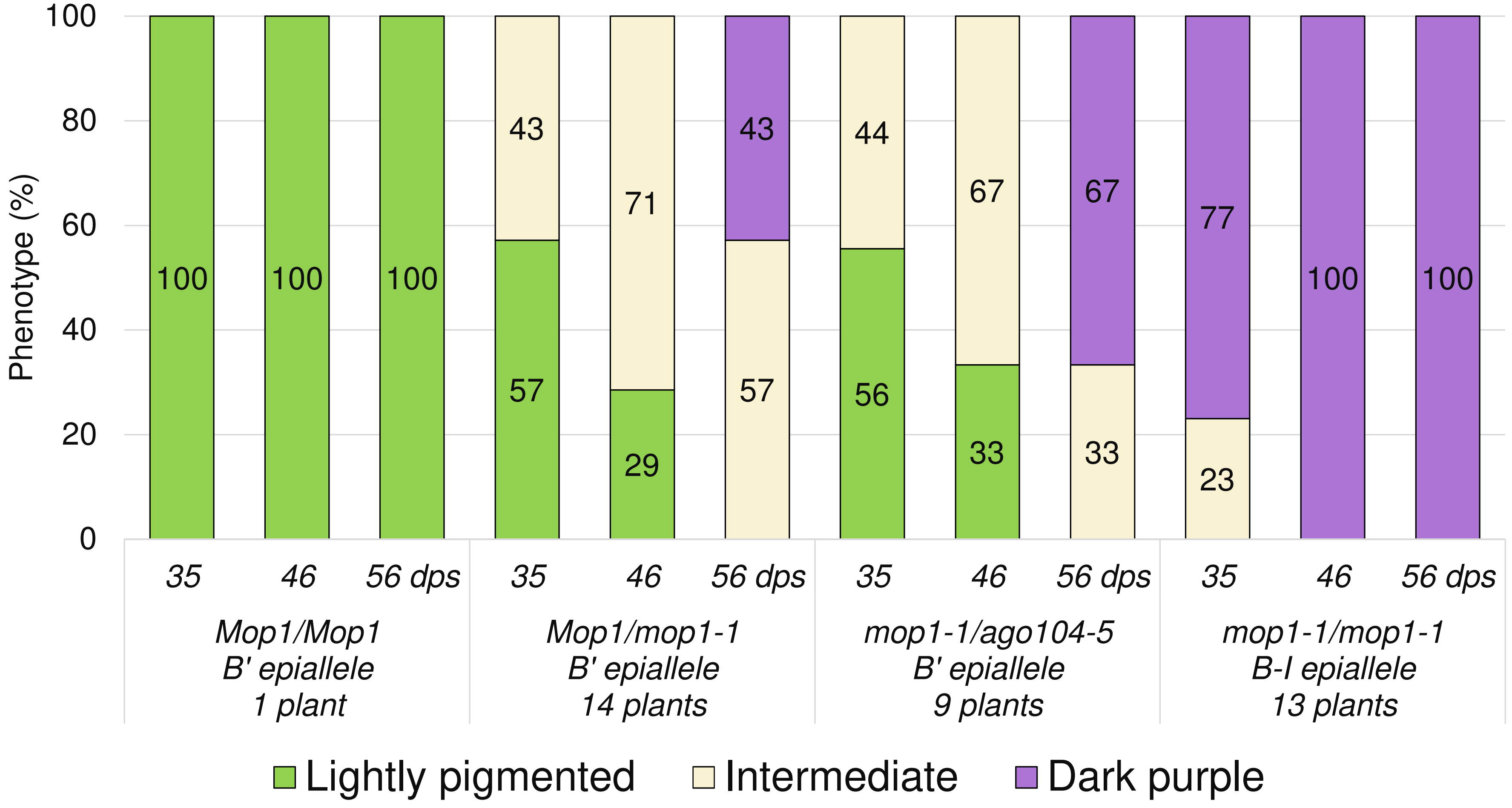


**S2 Fig. Evolution of mutant phenotypes at 35, 46 and 56 days post-seeding (dps).**

Mop1/Mop1 was described in Dorweiler et al. (2000). *mop1-1/ago104-5* plants are either double heterozygous or homozygous for the *ago104-5* mutation. *Mop1/mop1-1* plants inherited the non-mutant allele of *Ago104* in the last backcross of the paramutagenic population. The number of plants evaluated for each genotype is indicated in the horizontal axis title. Numbers in boxes are the percentage of occurrence of each phenotype.

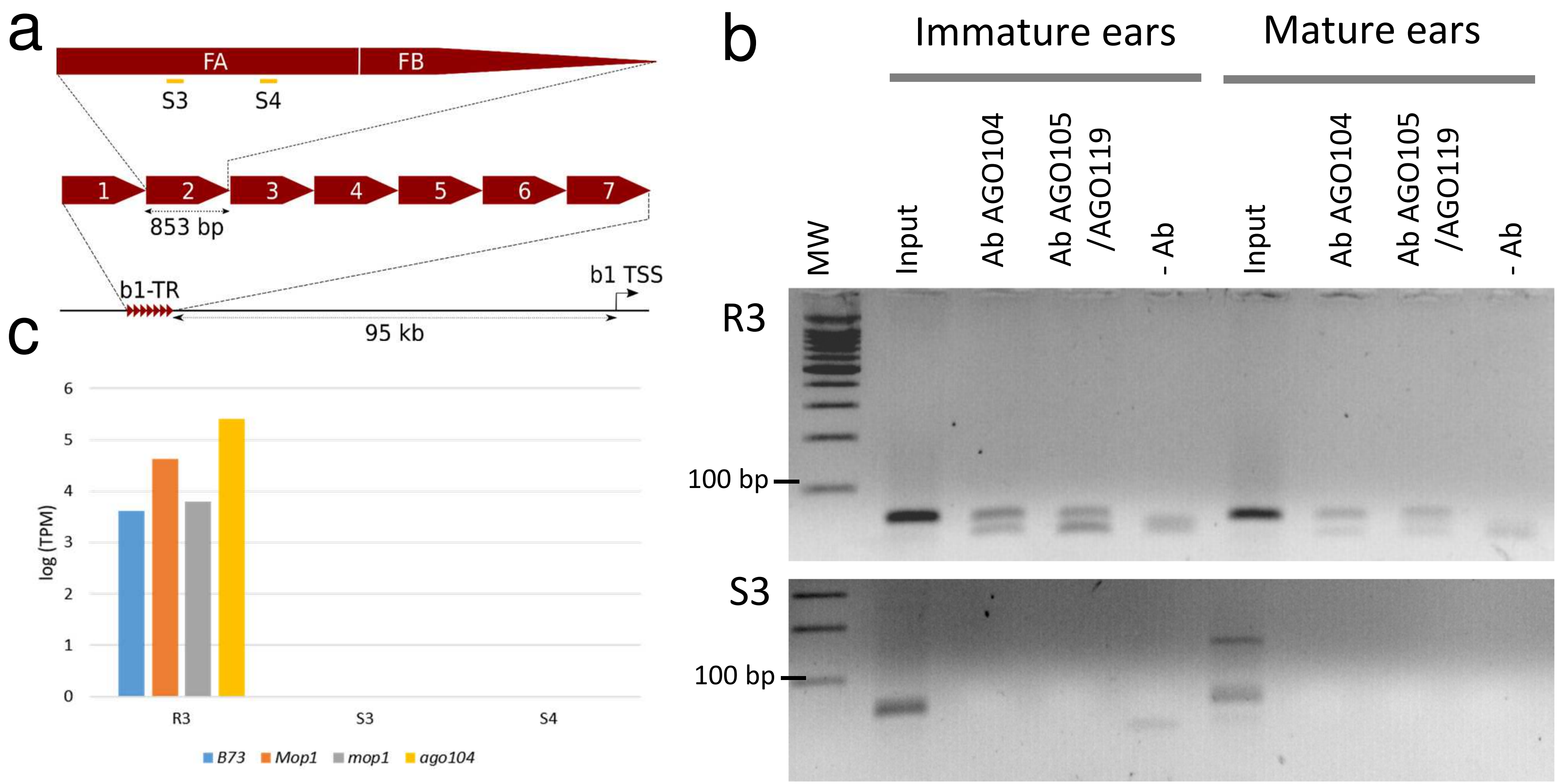


**S3 Fig. Presence of R3, S3 and S4 siRNAs loaded into AGO104 and AGO105/AGO119 in various genetic backgrounds.**

(a) Scheme of S3 and S4 location in the *b1TR*. FA and FB are the two halves of each repeat, as proposed by (Belele et al., 2013). Each repeat is 853 bp long, and they are separated from the *b1* transcription start site by 95 kb. (b) Stem-loop RT PCR of siRNAs extracted from IPs of AGO104 and of AGO105/AGO119. MW is the ladder. Input are small RNA extracted directly from the reproductive tissues. AbAGO104 and AbAGO105/AGO119 are the small RNA extracted from the IPs of AGO104 and AGO105/AGO119 respectively. -Ab correspond to the mock immunoprecipitation samples (without Ab). (c) Normalized expression levels of the sRNAs extracted from AGO104 in immature ears. Two technical repeats were sequenced for each genetic background (*B73*, *mop1-1* heterozygous, *mop1-1* homozygous and *ago104-5* homozygous mutants).

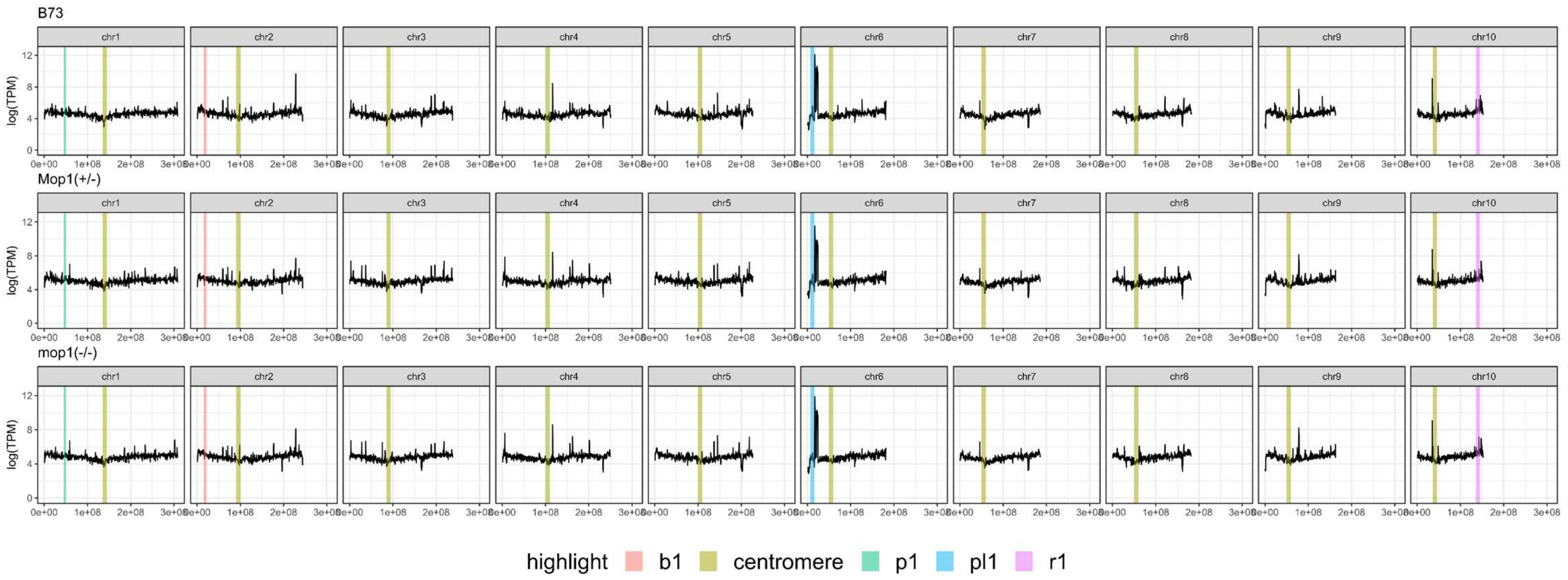


**S4 Fig. siRNA coverage on the 10 maize chromosomes of B73 reference genome (version 5).**
siRNAs were extracted from AGO104 IPs in immature ears of three genetic backgrounds (B73, *Mop1-1* heterozygous and *mop1-1* homozygous mutants) with 2 technical and biological repeats. Colored highlights are the positions of the centromeres and the four known paramutation loci in maize (*p1* on chromosome 1, *b1* on chromosome 2, *pl1* on chromosome 6 and *r1* on chromosome 10).

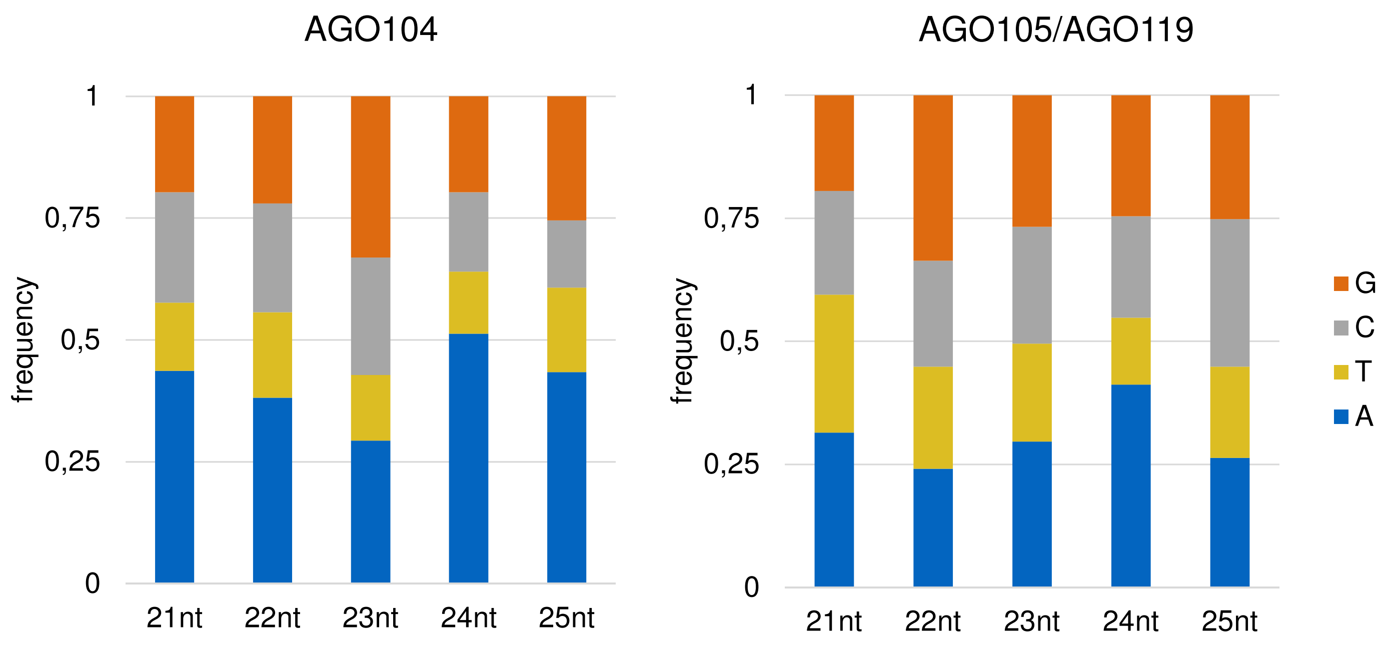


**S5 Fig. First base nucleotide bias in siRNAs bound by AGO104 and AGO105/AGO119.**

| Target protein | Target peptide | Peptide position | Organism |
|---|---|---|---|
| AGO104 | SERICKEQTFPLRQR | 326-341 | Rabbit |
| AGO105/119 | VGQFIKFDEMSETS | 853-867 | Rabbit |

**S1 Table. Antibodies characteristics**.

| Name | siRNA sequence | Size (nt) | Stem loop primer | TM | Forward primer | TM |
|---|---|---|---|---|---|---|
| R3 | ATGATTTGTGGGTCCGATGGCATA | 24 | GTCGTATCCAGTGCAGGGTCCGAGGTATTCGCACTGGATACGATATGCC | 73 | TGCCGATGATTTGTGGGTCC | 59 |
| S3 | GTTTGGAGACGATGACTCGTGGAC | 24 | GTCGTATCCAGTGCAGGGTCCGAGGTATTCGCACTGGATACGAGTCCAC | 75 | CCGGCGTTTGGAGACGATGA | 61 |
| S4 | GTTCAGTTCGTGGTGGACCGATGG | 24 | GTCGTATCCAGTGCAGGGTCCGAGGTATTCGCACTGGATACGACCATCG | 75 | GGGACGTTCAGTTCGTGGTG | 59 |

**S2 Table. Primer sequences used for siRNA stem loop RT-PCR.**
